# Connecting the sequence-space of bacterial signaling proteins to phenotypes using coevolutionary landscapes

**DOI:** 10.1101/044586

**Authors:** R. R. Cheng, O. Nordesjö, R. L. Hayes, H. Levine, S. C. Flores, J. N. Onuchic, F. Morcos

**Affiliations:** Center for Theoretical Biological Physics, Rice University, Houston, USA.; Department of Cell and Molecular Biology, Uppsala University, Uppsala, Sweden.; Department of Biophysics, University of Michigan, Ann Arbor, USA.; Department of Bioengineering, Rice University, Houston, USA.; Departments of Physics & Astronomy, Chemistry, and Biosciences, Rice University, Houston, USA.; Department of Biological Sciences, University of Texas at Dallas, Dallas, USA.

## Abstract

Two-component signaling (TCS) is the primary means by which bacteria sense and respond to the environment. TCS involves two partner proteins working in tandem, which interact to perform cellular functions while limiting interactions with non-partners (i.e., “cross-talk”). We construct a Potts model for TCS that can quantitatively predict how mutating amino acid identities affect the interaction between TCS partners and non-partners. The parameters of this model are inferred directly from protein sequence data. This approach drastically reduces the computational complexity of exploring the sequence-space of TCS proteins. As a stringent test, we compare its predictions to a recent comprehensive mutational study, which characterized the functionality of 20^4^ mutational variants of the PhoQ kinase in *Escherichia coli*. We find that our best predictions accurately reproduce the amino acid combinations found in experiment, which enable functional signaling with its partner PhoP. These predictions demonstrate the evolutionary pressure to preserve the interaction between TCS partners as well as prevent unwanted “crosstalk”. Further, we calculate the mutational change in the binding affinity between PhoQ and PhoP, providing an estimate to the amount of destabilization needed to disrupt TCS.

## Introduction

Early theoretical work on protein folding postulated that proteins have evolved to be minimally frustrated (Bryngelson and Wolynes 1987; Bryngelson, et al. 1995; Onuchic, et al. 1997), i.e., evolved to have favorable residue-residue interactions that facilitate folding into the native state while having minimal non-native energetic traps. The principle of minimal frustration provides intuition as to why protein sequences are not random strings of amino acids. The evolutionary constraint to fold into a particular, stable three-dimensional structure while minimizing the number of frustrated interactions greatly restricts the sequence-space of a protein (Leopold, et al. 1992; Bryngelson, et al. 1995; Onuchic, et al. 1997). Satisfaction of these constraints result in correlated amino acid identities within the sequences of a protein family. These correlated identities occur between different positions in a protein such as, for example, native contacts (Gobel, et al. 1994; Neher 1994; Shindyalov, et al. 1994). We refer to these quantifiable amino acid correlations as coevolution.

Of course, coevolution does not only arise from the constraint to fold. Proteins also fulfill cellular functions, which act as additional constraints on the sequences of proteins (Ferreiro, et al. 2014; Sikosek and Chan 2014; Wolynes 2015). In the context of signal transduction, proteins have evolved to be able to preferentially bind to a signaling partner(s) as well as catalyze the chemical reactions associated with signal transfer. An important example is two-component signaling (TCS) (Hoch 2000; Stock, et al. 2000; Laub and Goulian 2007; Casino, et al. 2010; Szurmant and Hoch 2010; Capra and Laub 2012), which serves as the primary means for bacteria to sense the environment and carry out appropriate responses. TCS consists of two partner proteins working in tandem: a histidine kinase (HK) and a response regulator (RR). Upon the detection of stimulus by an extracellular sensory domain, the HK generates a signal via autophosphorylation. Its RR partner can then transiently bind to the HK and receive the signal (i.e., phosphoryl group), thereby activating its function as a transcription factor. The HK has also evolved to catalyze the reverse signal transfer reaction (i.e., phosphatase activity), acting as a sensitive switch to turn off signal transduction. To prevent signal transfer with the wrong partner (i.e., “cross-talk”), TCS partners have mutually evolved amino acids at their respective binding interfaces that confer interaction specificity (Laub and Goulian 2007; Szurmant and Hoch 2010; Capra and Laub 2012). Thus, the collection of protein sequences of TCS partners contains quantifiable coevolution between the HK and RR sequences.

In principle, the determinants of interaction specificity for TCS can be quantified by a probabilistic model of sequence selection (Cheng, et al. 2014). Assuming that nature has sufficiently sampled the sequence-space of TCS proteins, the collection of protein sequences of TCS partners can be viewed as being selected under quasi-equilibrium from a Boltzmann distribution:

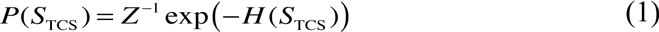

where *S*_TCS_ is the concatenated amino acid sequence of a HK and RR protein, *P* is the probability of selecting *S*_TCS_, and -*H* is proportional to the additive fitness landscape that governs the evolutionary sequence selection for TCS partners. Specifically, Eq. 1 was previously derived using simple models of evolutionary biology (Sella and Hirsh 2005), where *H* = −*vx* and *v* is the population size of the genotypes and *x* is the additive fitness (i.e., the negative log of the fitness); See Materials and Methods for more details. *H* is referred to in our work as a coevolutionary landscape.

Recently, maximum entropy-based approaches referred to as Direct Coupling Analysis (DCA) (Weigt, et al. 2009; Morcos, et al. 2011; Ekeberg, et al. 2013) have been successfully applied to infer the parameters of *H* (a Potts model) that governs the empirical amino acid sequence statistics. This has allowed for the direct quantification of the coevolution in protein sequence data (See Review: (de Juan, et al. 2013)). Early work using DCA to study TCS primarily focused on identifying the key coevolving residues between the HK and RR (Weigt, et al. 2009). Highly coevolving residue pairs have been used as docking constraints in a molecular dynamics simulation to predict the HK/RR signaling complex (Schug, et al. 2009), the autophosphorylation structure of a HK (Dago, et al. 2012), and the homodimeric form (transcription factor) of the RR (dos Santos, et al. 2015). DCA has also been applied to quantify the determinants of interaction specificity between TCS proteins (Procaccini, et al. 2011; Cheng, et al. 2014), building on earlier coevolutionary approaches (Li, et al. 2003; Burger and van Nimwegen 2008). In particular, DCA was used to predict the effect of point mutations on TCS phosphotransfer *in vitro* as well as demonstrate the reduced specificity between HK and RR domains in hybrid TCS proteins (Cheng, et al. 2014).

The experimental effort to determine the molecular origin of interaction specificity in TCS proteins (See Reviews: (Laub and Goulian 2007; Casino, et al. 2010; Szurmant and Hoch 2010; Podgornaia and Laub 2013)) precedes the recent computational efforts. Full knowledge of the binding interface between HK and RR was made possible through X-ray crystallography (Casino, et al. 2009). Scanning mutagenesis studies (Tzeng and Hoch 1997; Qin, et al. 2003; Capra, et al. 2010) provided insight on the subset of important interfacial residues that determine specificity. These key residues were mutated to enable a TCS protein to preferentially interact with a non-partner *in vitro* (Skerker, et al. 2008; Capra, et al. 2010). However, the extent of possible amino acid identities that allow TCS partners to preferentially interact *in vivo* has remained elusive until recent comprehensive work by Podgornaia and Laub (Podgornaia and Laub 2015). Their work focused on the PhoQ/PhoP TCS partners in *E. coli*, which control the response to low magnesium stress. PhoQ (HK) phosphorylates and dephosphorylates PhoP (RR) under low and high magnesium concentrations, respectively. Using exhaustive mutagenesis of 4 residues of PhoQ (20^4^ = 160,000 mutational variants) at positions that form the binding interface with PhoP, Podgornaia and Laub (Podgornaia and Laub 2015) were able to characterize all mutants based on their functionality in *E. coli*. It was found that roughly 1% of all PhoQ mutants were functional, enabling *E. coli* to exhibit comparable responses to magnesium concentrations as the wild type PhoQ. This finding uncovered a broad degeneracy in the sequence-space of the HK protein that still maintained signal transfer efficiency as well as interaction specificity with its partner.

We ask whether amino acid coevolution inferred using DCA could capture the functional mutational variants observed in the comprehensive mutational study of PhoQ and if so, to what extent? Capturing this functionality requires that information gleaned from coevolution is sufficient to estimate the effect of mutations to PhoQ on its interactions with PhoP as well as on unwanted “cross-talk”. Hence, our question is important to determine if coevolutionary methods can be extended from studying two interacting proteins to studying an interaction network (e.g., systems biology). Further, this question is of particular interest to those who want to engineer novel mutations in TCS proteins that can maintain or encode the interaction specificity of a TCS protein to its partner or a non-partner, respectively.

To answer this question, we first infer a Potts model, *H* (see Eq. 1), which forms the basis for quantifying how mutations affect the interaction between a HK and RR protein. Focusing on the parameters of *H* that are related to interprotein coevolution, we construct a coevolutionary landscape to quantify TCS interactions, *H*_TCS_, for a given sequence of an HK and RR protein. *H*_TCS_ serves as a proxy for signal transfer efficiency, allowing us to quantify the effect on fitness of the interaction between any HK and RR protein. Further, we can assess how mutations affect fitness due to changes in the HK/RR interaction by computing the mutational change in *H*_TCS_ between the mutant sequence, 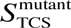, and the wild type sequence, 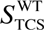:

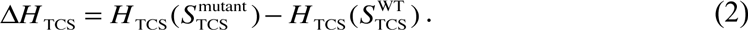

Considering the concatenated sequence of PhoQ and PhoP, we compute Eq. 2 for the 20^4^ PhoQ mutational variants. We find that mutants with the most favorable Δ*H*_TCS_ (e.g., most negative) were classified as functional HKs by Podgornaia and Laub (Podgornaia and Laub 2015)—i.e., true positive predictions. Next, we focus on mutations predicted to be favorable by Eq. 2 that were classified as non-functional in experiment. Expanding our analysis of the PhoQ mutants beyond its interaction with PhoP, we consider how mutations affect the signal transfer efficiency, *H*_TCS_, between PhoQ and all of the RR proteins in *E. coli*. We find that many of these nonfunctional mutants exhibit “cross-talk” interactions according to our model, accounting for their non-functionality. If we exclude these promiscuous variants, we can better isolate the true positive predictions that are functional from false positives that are non-functional. Our predictions also capture context-dependent mutational affects that were observed in experiment, i.e., epistasis. Finally, we estimate the mutational change in binding affinity in the PhoQ/PhoP bound complex using the Zone Equilibration of Mutants (ZEMU) method (Dourado and Flores 2014), a combined physics- and knowledge-based approach for free energy calculations. Consistent with what we would expect, we find that mutations that destabilize the HK/RR interaction tend to be non-functional with very high statistical significance. Non-functional mutants are on average destabilized by ~2 kcal/mol with respect to functional mutants.

The work described herein demonstrates that a coevolutionary model (i.e., additive fitness landscape) built from sequence data can directly connect molecular details at the residue-level to mutational phenotypes in bacteria. This has broad applications in systems biology, but also in synthetic biology since our computational framework can be used to select mutations that enhance or suppress interactions between TCS proteins. A more detailed description of our computational approaches can be found in the Materials and Methods section.

## Results

### *Mutational change in coevolutionary landscape, Δ*H*_TCS_, for PhoQ/PhoP interaction*

We first focus on the parameters of the inferred Potts model (Eq. 3) that describe the coevolution between the Dimerization and Histidine phosphotransfer domain (DHp) and the Receiver (REC) domain (Fig. 1A), which form the HK/RR binding interface (Fig. 1B). The interprotein statistical couplings (Fig. 1C) of Eq. 3 are used to construct a coevolutionary landscape, *H*_TCS_ (Eq. 5), as a proxy for signal transfer efficiency. For each of the 1,659 functional and 158,341 non-functional PhoQ-mutational variants identified by Podgornaia and Laub (Podgornaia and Laub 2015), we compute the mutational change, Δ*H*_TCS_, between PhoQ and PhoP. As an initial step, we only consider the PhoQ/PhoP sequence, i.e., we do not yet consider other RR proteins than PhoP. A histogram of Δ*H*_TCS_ is generated for all mutational variants (Fig. 2A). The distribution of the functional mutants tends more towards favorable Δ*H*_TCS_ than the distribution of non-functional mutants, but more interestingly, the most favorable predictions of our model contain mostly functional mutations. This is made clear by a plot of the Positive Predictive Value (PPV) for the top *N* mutational variants ranked by Δ*H*_TCS_ (Fig. 2B) from most favorable to most deleterious. The top 25 mutational variants ranked by Δ*H*_TCS_ contain 20 functional mutants and 5 nonfunctional mutants (i.e., PPV=0.8).

**Fig. 1.**
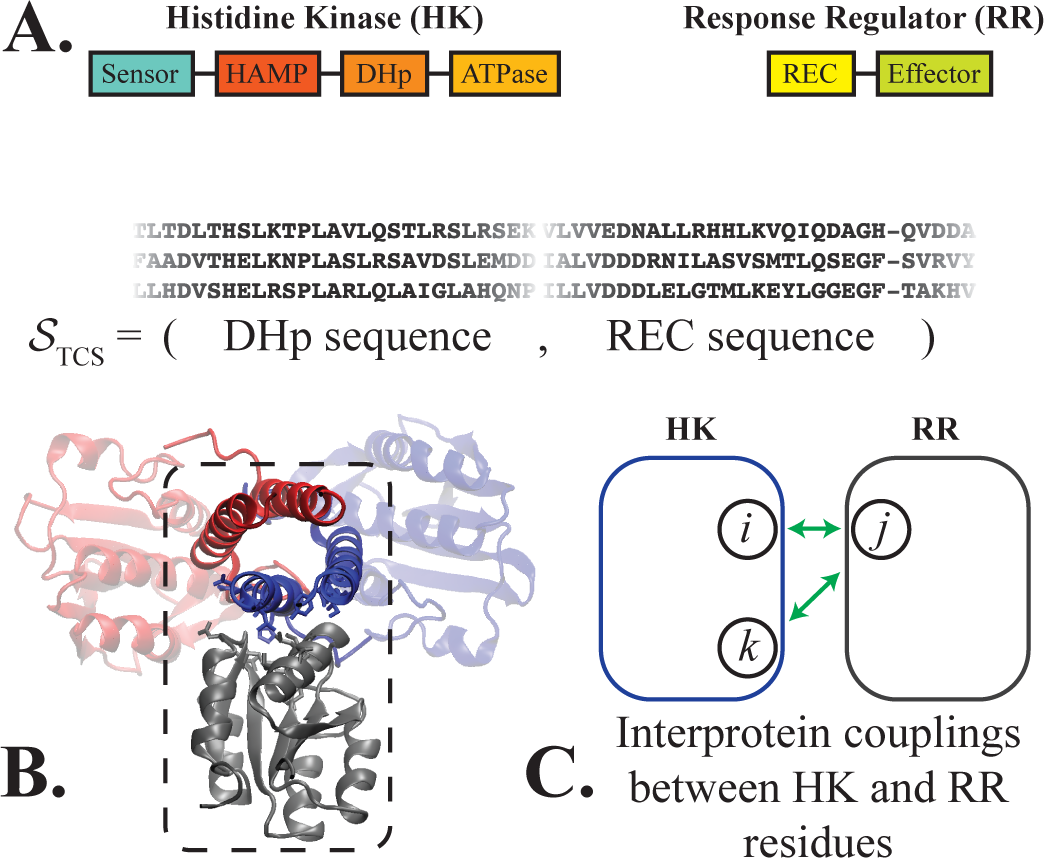
TCS domain interactions of interest. We focus only on HK proteins that have the following domain architecture from N to C terminus: sensor, HAMP, DHp, and ATPase. Likewise, we consider RR proteins that consist of a REC domain followed by an effector domain. (A) The interaction between the DHp and REC domains of the HK and RR proteins, respectively, form the TCS complex. Sequences of TCS partners are collected and stored as the concatenated sequence of the DHp and REC domains, *S*_TCS_ (See Materials and Methods). (B) A representative structure of the HK/RR TCS complex previously predicted for the KinA/Spo0F complex in *B. subtilis (Cheng, et al. 2014)*. The HK homodimer is shown in red and blue while the receiver domain of the RR is shown in gray. The dashed box highlights the DHp and REC interface. This predicted complex is consistent with the experimentally determined crystal structure of HK853/RR468 of *T. maritima* (Casino, et al. 2009) as well as another computationally predicted TCS complex (Schug, et al. 2009). (C) Our proxy for signal transfer efficiency, *H*_*TCS*_ (Eq. 5), is composed of the statistical coupling parameters that describe coevolution between interprotein residues (depicted in green). Hence, *H*_*TCS*_ naturally captures the context-dependence of mutating a residue in the HK when a residue in the RR is also mutated, or vice versa (See Materials and Methods for more details).

**Fig. 2.**
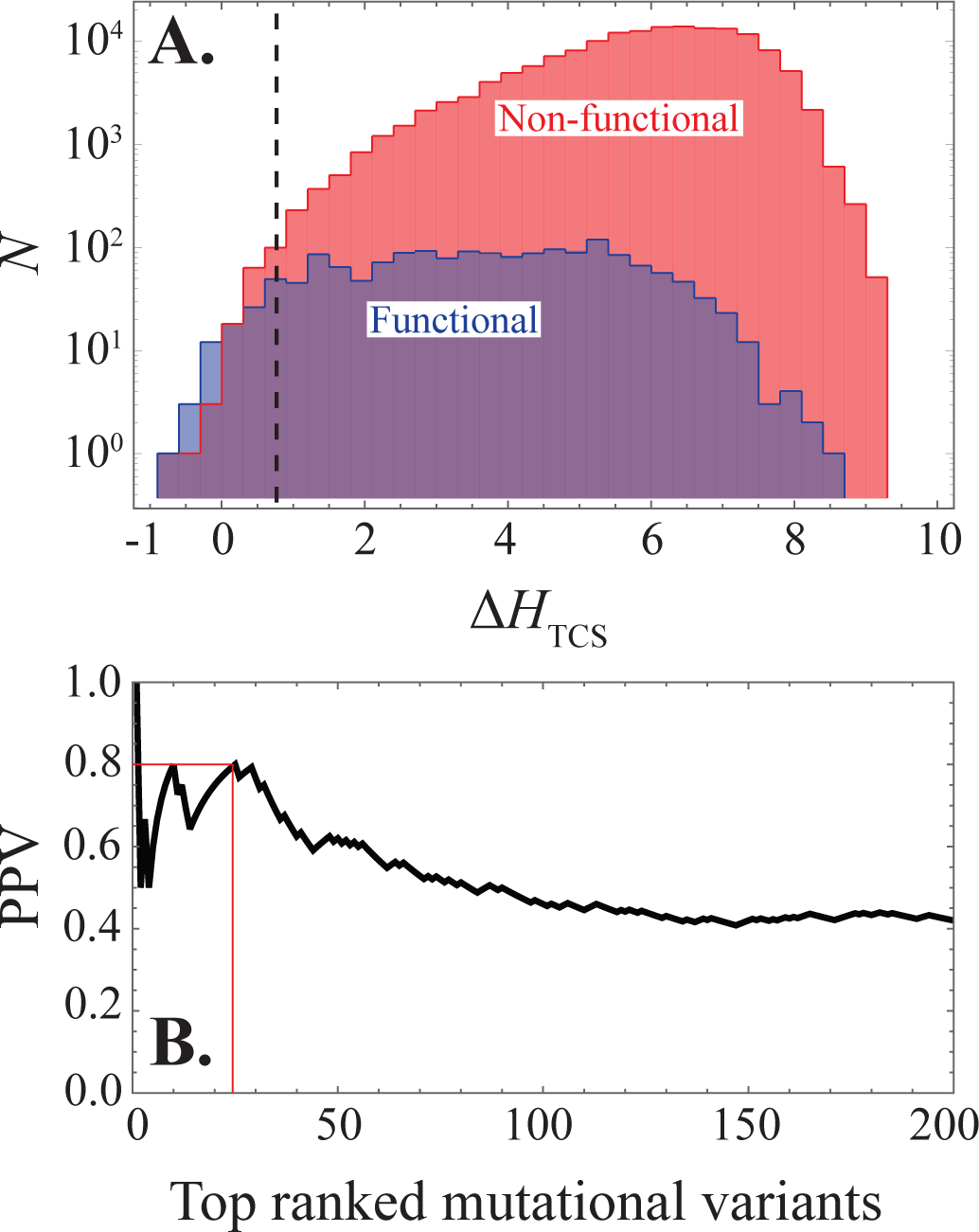
Effect of mutations on the PhoQ/PhoP interaction. (A) Considering the concatenated sequence of PhoQ/PhoP, a histogram of Δ*H*_TCS_ (Eq. 5) is plotted for the functional (blue) and non-functional (red) mutational variants reported by Podgornaia and Laub (Podgornaia and Laub 2013). The color purple shows parts of the plot where the blue and red histograms overlap. The dashed line roughly partitions the 200 most favorable mutational variants given by Δ*H*_TCS_, which contains more functional than nonfunctional mutants. By definition, Δ*H*_TCS_ = 0 corresponds to the wild type PhoQ/PhoP and Δ*H*_TCS_ = 0 corresponds to mutations that we predict to be more favorable to PhoQ/PhoP signaling than the wild type. (B) We plot the positive predictive value (PPV) as a function of the *N* mutational variants ranked by Δ*H*_TCS_ from the most to least favorable for the first 200 mutants. PPV=TP/(TP+FP), where true positives (TP) and false positives (FP) refer to the fraction of mutants that are functional or non-functional, respectively, in the top *N* ranked variants. The thin red lines denote that the top 25 ranked mutational variants have a PPV of 0.8.

### System-level analysis using ΔH_TCS_: functional mutants limit “cross-talk”

Mutations that may enhance signal transfer efficiency between PhoQ and PhoP *in vitro* may still result in a non-functional PhoQ/PhoP system *in vivo*. This would occur if the mutations to PhoQ sufficiently encoded it to preferentially interact with another RR in *E. coli*. For this reason, we focused our computational analysis on the subset of mutational variants that preserve PhoQ/PhoP specificity by limiting “cross-talk” according to our coevolutionary model.

We first calculate the proxy for signal transfer efficiency, *H*_TCS_, between the wild type PhoQ sequence and all of the non-hybrid RR proteins in *E. coli* (Fig. 3A). We find that for wild type PhoQ, the most favorable *H*_TCS_ (most negative) is with its known signaling partner, PhoP. As a consistency check, we also plot *H*_TCS_ for different combinations of the cognate partners TCS proteins in *E. coli* (Fig. S1). This result is consistent with previous computational predictions that used information-based quantities (Procaccini, et al. 2011; Cheng, et al. 2014) to quantify interaction specificity.

Extending upon Fig. 3A, we assess “cross-talk” in our model by calculating *H*_TCS_ between each PhoQ mutant and all of the non-hybrid RR in *E. coli*. We exclude all mutant-PhoQ variants that have a more favorable *H*_TCS_ with a non-partner RR. These excluded mutants are excellent candidates for engineering specificity in *E. coli*. Applying our exclusion criterion, we find that only 181 functional and 1,532 non-functional variants remain, i.e., 89% and 99% of the functional and non-functional variants, respectively, were removed. A histogram of the remaining (cross-talk excluded) mutants as a function of Δ*H*_TCS_ (Fig. 3B) shows that a filter based on interaction specificity is better able to isolate the true positive (functional) variants. Notably, the first 17 ranked variants are all functional variants. Once again, ranking the filtered variants by Δ*H*_TCS_ from the most favorable to the least favorable, we can plot the PPV (Fig. 3C) for the top *N* ranked variants. We find that the cross-talk excluded PPV tends to lie above the original PPV from Fig. 2B.

**Fig. 3.**
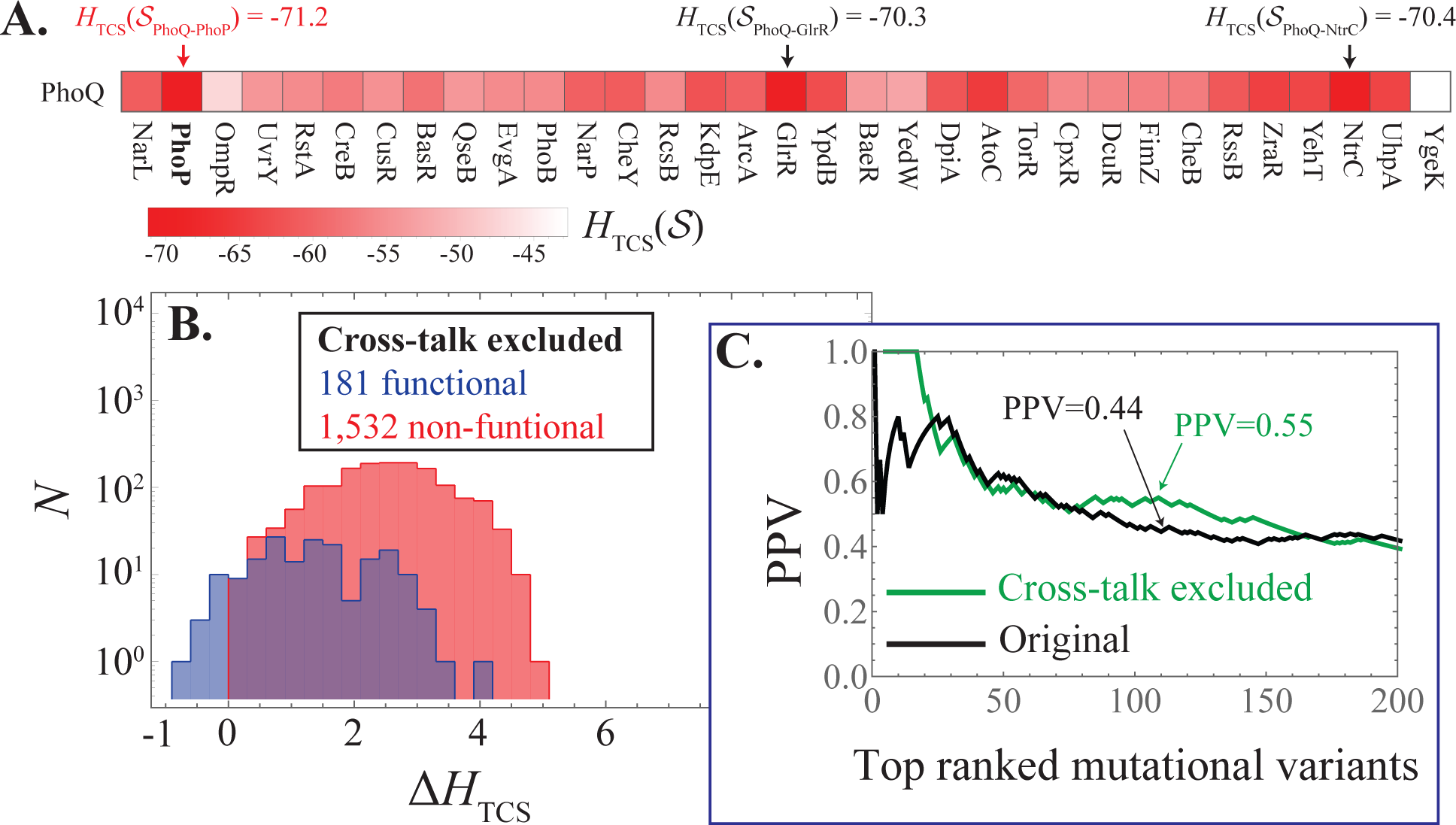
Excluding mutational variants that are inferred to “cross-talk”. (A) A grid plot showing *H*_TCS_ (Eq. 5) computed for the wild type PhoQ sequence with all of the non-hybrid RR protein sequences in *E. coli*, respectively. The most favorable interaction (most negative) given by *H*_TCS_ is between PhoQ and its partner PhoP. (B) We plot the Cross-talk excluded subset (181 functional 1,532 non-functional) in a histogram as a function of the Δ*H*_TCS_ similar to Fig. 2A. (C) We plot the PPV as a function of the *N* top mutational variants ranked by Δ*H*_TCS_ for the first 200 mutants. The PPV for the cross-talk excluded mutational variants from Fig. 3B are plotted in green while the original PPV (Fig. 2B) is shown in black.

**Fig. 4.**
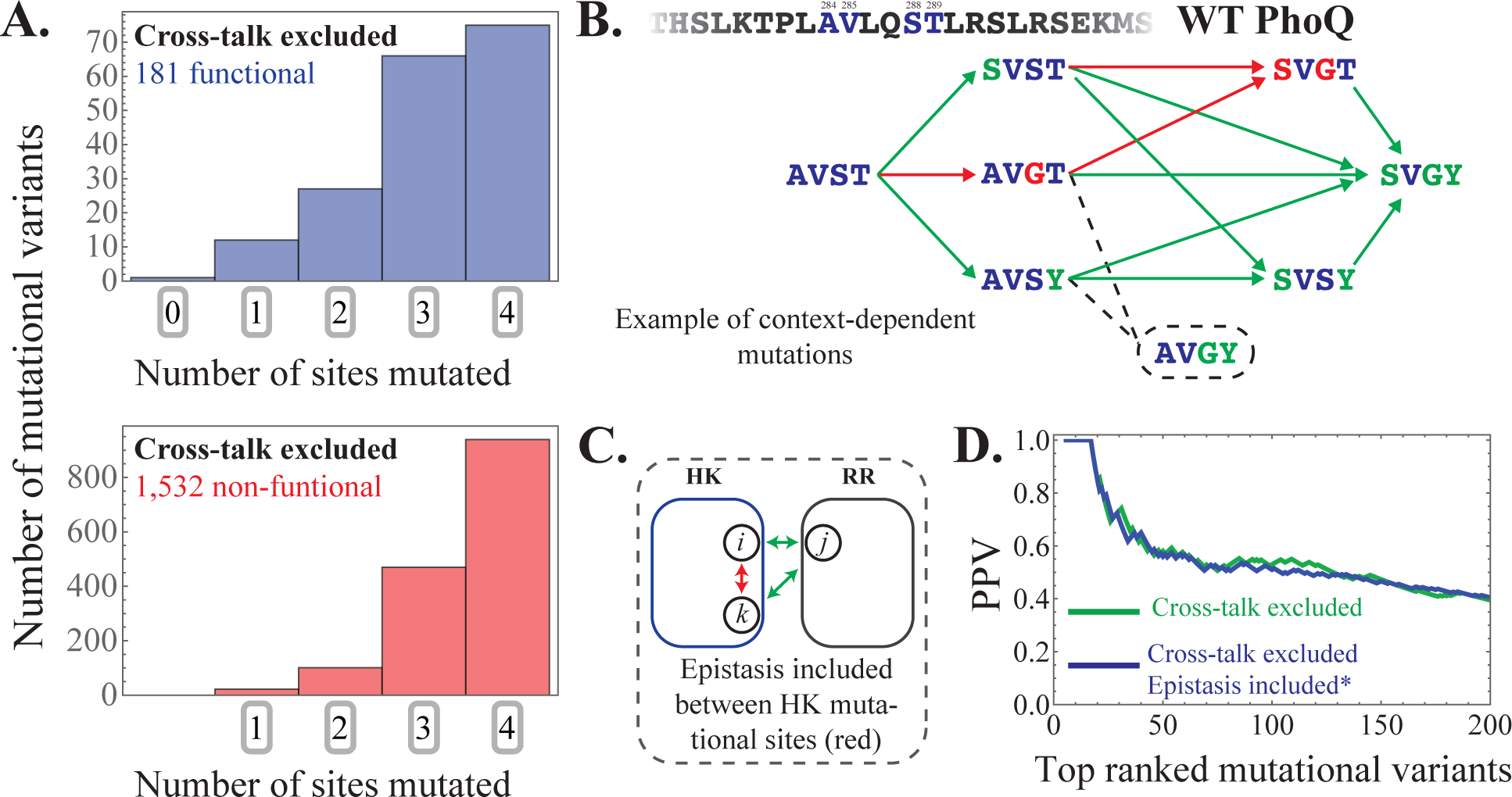
Capturing epistatic mutational effects. (A) Histograms of the 0, 1, 2, 3, and 4-site mutants are plotted for the 181 functional and 1,532 non-functional mutational variants in our cross-talk excluded subset that is predicted using *H*_TCS_ (Eq. 5). (B) An example of mutational context-dependence predicted by *H*_TCS_ is shown here, considering residues 254, 255, 258, 259 of PhoQ, which have the WT amino acid configuration AVST, respectively. Green arrows drawn from AVST indicate single point mutations that are correctly predicted to result in a functional phenotype. Successive green arrows indicate double and triple point mutations from the WT that are correctly predicted to be functional. Likewise, red arrows indicate single point mutations from an amino acid configuration that are correctly predicted to result in a non-functional phenotype. The AVGY mutation (dashed line and circle) was found to be functional in experiment but is predicted to cross-talk by *H*_TCS_. (C) The schematic shows intraprotein coevolution (red) between residues *i* and *k* (of the HK) and interprotein coevolution (green) between residues *i* and *j* as well as *k* and *j.* As previous described in Fig. 1C, interprotein coevolution captures the effects of mutating the HK when the RR is also mutated (or vice versa). Epistasis between HK and RR proteins are naturally incorporated within *H*_TCS_. On the other hand, epistasis between the 4-mutational sites of the Podgornaia and Laub experiment is described in our model through the statistical couplings between the 4-mutational sites (red in schematic). These additional parameters are added to *H*_TCS_ to obtain 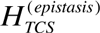 (See Materials and Methods). (D) We plot the PPV as a function of the *N* top mutational variants ranked by Δ*H*_TCS_ for the first 200 mutants. The PPV predicted by *H*_TCS_ (Fig. 3C) is shown in green while the PPV predicted by 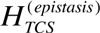 is shown in blue.

### Mutational context-dependence in the coevolutionary landscape

A significant finding of the Podgornaia and Laub study was the context-dependent nature of the many of the mutations. For example, individual mutations that may result in a non-functional phenotype may result in a functional phenotype when combined. It is well known that mutations may exhibit such a context dependence, or epistasis, i.e., mutations introduced together have an effect on fitness that is not simply the combined effect of each mutation alone. Such effects would restrict the connectivity between functional mutations and act as constraints on TCS evolution (Podgornaia and Laub 2015).

We find that the functional predictions in Fig. 3 tend to be 3- and 4-point mutations, highlighting their non-trivial nature (Fig. 4A). While the effect of mutating a HK protein when its partner RR is mutated (and vice versa) is naturally captured in our model through interprotein statistical couplings (Fig. 1C), the mutational context-dependence (i.e., intraprotein couplings) of HK only mutations or RR only mutations is not explicitly contained within Eq. 5. Even with these limitations, the model is still able to distinguish between functional multi-point mutations that are composed of non-functional single-point mutations for the most favorable predictions, in accordance with experiment (Podgornaia and Laub 2015). One example is provided in Fig. 4B for the mutation of WT PhoQ from AVST at residues 284, 285, 288 and 289, respectively, to SVGY (i.e., a 3-point mutation). For single point mutations from AVST, *H*_*TCS*_ correctly predicts that SVST and AVSY are functional while AVGT is non-functional. For two point mutations, *H*_*TCS*_ correctly predicts that SVSY is functional while SVGT is non-functional. More interestingly, *H*_*TCS*_ finds that when the non-functional 2-point mutation SVGT is combined with a Tyrosine mutation to position 289 (i.e., SVGY), the functionality is recovered. It should be noted that the two-point mutation AVGY is predicted to have a favorable Δ*H*_*TCS*_ between PhoQ and PhoP, but is predicted to undergo cross-talk, and thus, is discarded despite being functional in experiment.

We next consider a model where epistasis between the 4-mutational sites explored by Podgornaia and Laub are explicitly introduced into Eq. 5 (See Eq. 6 in Materials and Methods). The epistatic effects of multiple HK mutations are captured by the statistical couplings of the Potts model that describe the pairwise coevolution between the HK mutational sites. This is illustrated schematically in Fig. 4C. We find that such a model, 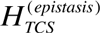 (Eq. 6), is consistent with the original model, *H*_*TCS*_, in terms of its predictions and predictive quality (Fig. 4D). In particular, the prediction of functional phenotypes for the example in Fig. 4B yields identical results. Nevertheless, this model provides additional epistatic mutational effects that are not simply the added sum of the individual mutational effects.

Finally, we examine a model that only considers HK coevolution in SI Main Text. This model naturally includes the epistasis between the 4-mutational sites explored in experiment (Podgornaia and Laub 2015). From this model, we find that coevolution between HK residues alone is insufficient for capturing the functional phenotypes observed in experiment.

### Mutational change in the binding affinity using a combined physics- and knowledge-based approach

We used ZEMu (See Materials and Methods) to compute the mutation-induced change in the binding affinity, 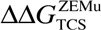, between PhoQ and PhoP. The calculation converged for 42,985 mutants (702 functional and 42,283 non-functional) from a randomly selected subset of the 20^4^ variants. We first examined a scatter plot of 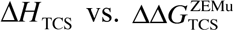 (Fig. S6) to observe whether a functional relationship can be deduced from the computational data alone. The mutational change in the coevolutionary landscape, Δ*H*_TCS_, would in principle exhibit a non-linear dependence on 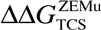, while also depending on many other quantities including the binding affinity with other proteins. We find no obvious relationship between the two quantities in our current analysis.

A histogram of 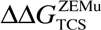 is plotted for the 42,985 mutants in Fig. 5A. A histogram of Δ*H*_TCS_ for the same subset of mutants is shown in Fig. S7A. On the population level, functional mutations exhibit a mean 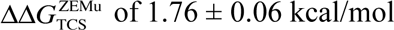 lower than that of the non-functional mutants, with a Wilcoxon rank-sum test p-value < 2.22×10^−16^. This indicates that destabilizing mutations of ~2 kcal/mol are sufficient for disrupting TCS. Furthermore, destabilizing mutations that are more than 2 standard deviations greater than the mean 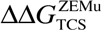 for functional variants are significantly less likely to be functional, with a p-value < 10^−6^ computed from a cumulative binomial distribution (based on the 6157 mutants above this threshold, 19 of which are functional).

We next examine the potentially deleterious effect of mutations that overly stabilize the binding affinity between PhoQ and PhoP. Although we find that all 56 mutants with 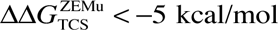 are non-functional (Fig. 5A), this has no statistical significance (p-value ~ 0.4). Fig. 5A illustrates our point that the functional mutations tend to have a neutral affect on 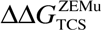, while high 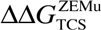 is strongly associated with the loss of function. Although very low 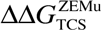 may visually appear to be enriched with non-functional mutants, this is based on a small number of mutants and does not have statistical significance.

**Figure 5.**
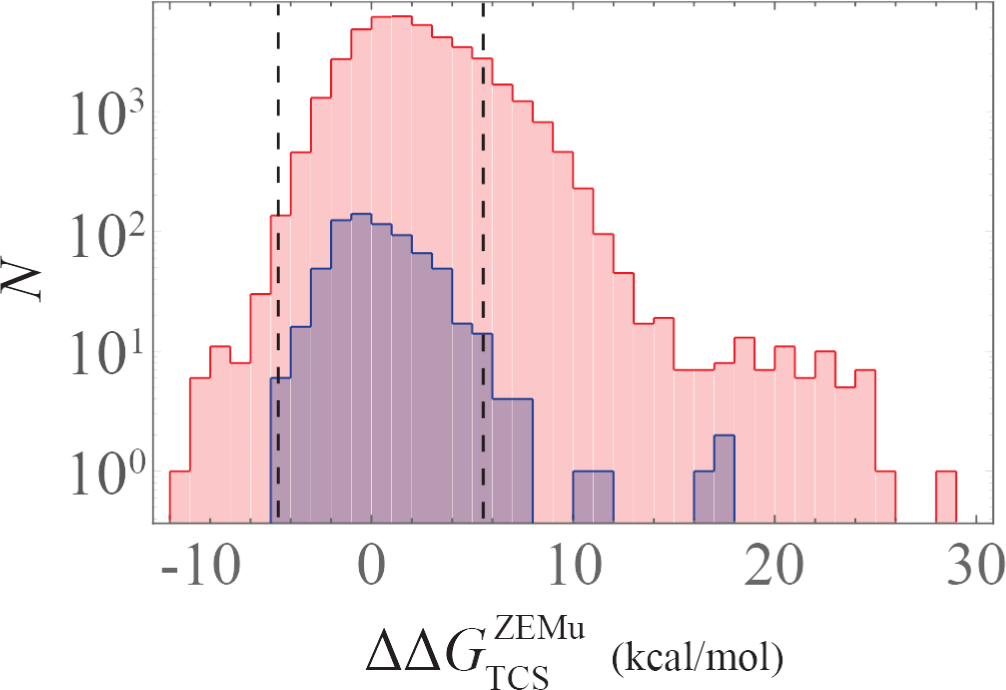
Mutational change in binding affinity for PhoQ/PhoP interaction. A histogram of 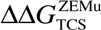 (See Materials and Methods), is plotted for the 702 functional (blue) and 42,283 non-functional (red) mutational variants analyzed in our study. The dashed lines denote ±2 standard deviations from the mean of the functional (blue) distribution.

## Discussion

Treating a large collection of amino acid sequence data for TCS partner proteins as independent samples from a Boltzmann equilibrium distribution, we infer a coevolutionary landscape, *H*_TCS_. Specifically, *H*_TCS_ is proportional to the negative of the additive fitness landscape, which captures the coevolving amino acid combinations that give rise to interaction specificity in TCS systems. In the past, we were able to predict how a point mutation to a TCS protein affects its ability to transfer signal to its partner *in vitro* (Cheng, et al. 2014). Our present work shifts the paradigm of coevolution-based analysis towards systems biology by extending our analysis to include how those mutations affect “cross-talk” in a bacterial organism. We demonstrate that our most favorable predictions for multiple site mutations can accurately capture *in vivo* TCS functionality, consistent with the comprehensive mutational study of Podgornaia and Laub (Podgornaia and Laub 2015). This is not a trivial computational task, since inferring coevolutionary information from sequence data is highly underdetermined and estimates ~10^7^ parameters (for Eq. 3) from ~10^3^ sequences of TCS partners. Adding to the problem complexity, it is plausible that the full functional sequence space has not yet being explored by evolutionary process (Capra and Laub 2012; Echave, et al. 2016). Despite this, the coevolutionary landscape is predictive and identifies mutational variants that are not found in nature, e.g., none of the mutational sequences are included as input data in our model. We have demonstrated the feasibility of generating predictions using coevolution and the predictive power of such an approach will only systematically improve as more sequences of TCS partners are collected.

Similar predictions to those discussed herein can readily be used to engineer novel protein-protein interactions in TCS systems. Such a strategy would potentially complement already existing strategies to match novel inputs with outputs via modular engineering (Tabor, et al. 2011; Whitaker, et al. 2012; Ganesh, et al. 2013; Schmid, et al. 2014; Hansen and Benenson 2016). The strength of our coevolutionary approach is that it makes possible an efficient search of sequence-space for mutations at arbitrary positions in either the HK or RR that desirably enhance or suppress its interaction with a RR or HK, respectively. It can also readily be applied to study the *in vivo*, system-level effect of mutating a TCS protein on insulating its interaction with a desired partner or enabling “cross-talk” with non-partners. Our study highlights an intuitive but key principle for selecting mutations to a TCS protein that encodes specificity *in vivo*: mutations must be selected to enhance protein-protein interactions with a desired partner while limiting protein-protein interactions with undesired partners. While also intuitive, we demonstrate that mutations that significantly destabilize the binding affinity result in the loss of signaling. Further, we estimate that destabilization of ~2 kcal/mol in the binding affinity between TCS partners is sufficient to disrupt TCS.

It is also important to note that coevolutionary methods described here for identifying mutational phenotypes (e.g., response to magnesium stress) is that they are not particular to TCS systems. This framework is transferable to other systems where molecular interactions coevolve to preserve function, opening the window to a large set of open problems in molecular and systems biology. Our results further extend the idea that a combination of coevolutionary based methods, molecular modeling and experiment can be used to identify the proper amino acids sites and identities that can be used to identify mutational phenotypes. Our study highlights the important role of coevolution in maintaining protein-protein interactions, as in the case of bacteria signal transduction. Statistical methods that probe coevolution not only allow us to connect molecular, residue-level details to mutational phenotypes, but also to explore the evolutionary selection mechanisms that are employed by nature to maintain interaction specificity, e.g., negative selection (Zarrinpar, et al. 2003). Further investigations of other systems that are evolutionarily constrained to maintain protein-protein interactions could elucidate the extent at which our methods can be used in alternative systems. One potential example is the toxin-antitoxin protein pairs in bacteria, which was the focus of recent experimental work (Aakre, et al. 2015) elucidating the determinants of interaction specificity.

## Materials and Methods

### Sequence database for HK and RR inter-protein interactions: DHp and REC

We obtain multiple sequence alignments (MSA) from Pfam (Finn, et al. 2014) (Version 28), focusing on the DHp (PF00512) and REC (PF00072) domains of the HK and RR, respectively (Fig. 1A). The first 4 positions (columns) of PF00512 were removed due to poor alignment of the PhoQ sequence at those positions. The remaining DHp MSA has a length of *L*_DHp_ = 60. Each REC MSA had a default length of *L*_REC_ = 112. Here, we considered HK proteins that have the same domain architecture as the PhoQ kinase from *E. coli*, i.e., DHp domain sandwiched between an N-terminal HAMP domain (PF00672) and a C-terminal ATPase domain (PF02518). The remaining HK (DHp) sequences were paired with a TCS partner RR (REC) by taking advantage of the observation that TCS partners are typically encoded adjacent to one another under the same operon (Skerker, et al. 2005; Yamamoto, et al. 2005), i.e., ordered locus numbers differ by 1. Further, we exclude all TCS pairs that are encoded adjacent to multiple HKs or RRs. Each DHp and REC sequence that was paired in this fashion was concatenated into a sequence (Fig. 1A), *S*_TCS_ = (*A*_1_, *A* _2_, …, *A*_*L*−1_, *A*_*L*_) of total length *L* where *A*_*i*_ is the amino acid at position *i* which is indexed from 1 to *q* = 21 for the 20 amino acids and MSA gap. The DHp sequence is indexed from positions 1 to *L*_DHp_ and REC sequence from positions *L*_DHp_ = 1 to the total length of *L* = *L*_DHp_ + *L*_REC_ = 172. Our remaining dataset consisted of 6,519 non-redundant concatenated sequences.

### Inference of parameters of coevolutionary model

An amino acid sequence *s* = (*A*_1_, *A* _2_, …, *A*_*L*_) for a protein or interacting proteins can be viewed as being selected from a Boltzmann equilibrium distribution, i.e., *P*(*s*) = *Z*^−1^ exp(−*H*(*s*)). The Boltzmann form of *P* was previously derived for an evolving population in the limit where the product of the population size and mutation rate is very small (Sella and Hirsh 2005). Specifically, it was shown (Sella and Hirsh 2005) that *H* (*s*) = −*vx*(*s*), where *v* is the population size and *x* (*s*) is the additive fitness landscape (i.e., log of the fitness). A high population size suggests many viable sequences that a protein can mutate to, which makes the population more robust to deleterious mutations. Related work (Halpern and Bruno 1998) modeled site-specific selection of sequences, which has been extended upon by numerous works (Tamuri, et al. 2012; Spielman and Wilke 2015; Bloom 2016).

Under certain limiting conditions, *H*(*s*) appears to share a correspondence with the energetics of protein folding (See review: (Sikosek and Chan 2014)). Assuming that the sequence diversity is completely due to stability considerations, *H*(*s* = *βE*(*s*) where *E*(*s*) is the energy of the folded protein with respect to the unfolded state and β = *k*_*B*_*T*_sel_)^−1^ is the inverse of the evolutionary selection temperature from protein folding theory (Pande, et al. 1997, 2000; Morcos, et al. 2014). Several studies have reported strong linear correlation between mutational changes in *H*(*s*) with mutational changes in protein stability (Lui and Tiana 2013; Morcos, et al. 2014; Contini and Tiana 2015). However, *H*(*s* = *βE*(*s*) may not be an appropriate approximation for proteins that have evolved with interacting partners, for which sequence selection is plausibly influenced by additional factors such as binding affinities as well as binding/unbinding rates.

Often it is of interest to solve the inverse problem of inferring an appropriate *H* (*s*) when provided with an abundant number of protein sequences. Typical approaches to this problem have applied the principle of maximum entropy to infer a least biased model that is consistent with the input sequence data (Weigt, et al. 2009; Morcos, et al. 2011), e.g., the empirical single-site and pairwise amino acid probabilities, *P*_*i*_(*A*_*i*_) and *P*_*ij*_ (*A*_*i*_, *A*_*j*_), respectively. The solution of which is the Potts model:

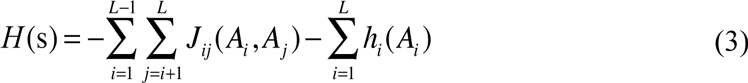

where *A*_*i*_ is the amino acid at position *i* for a sequence in the MSA, *J*_*ij*_ (*A*_*i*_, *A*_*j*_) is the pairwise statistical couplings between positions *i* and *j* in the MSA with amino acids *A*_*i*_ and *A*_*j*_, respectively, and *h*_*i*_ (*A*_*i*_) is the local field for position *i*. We estimate the parameters of the Potts model, {**J**, **h**}, using the pseudo-likelihood maximization Direct Coupling Analysis (plmDCA) (See Ref: (Ekeberg, et al. 2013) for full computational details). It is important to note that the mutational context-dependence (epistasis) between residues *i* and *j* is naturally captured in the model through the statistical couplings, *J*_*ij*_ (*A*_*i*_, *A*_*j*_).

Previous studies have applied DCA to a number of problems in structural biology. DCA has been used to identify highly coevolving pairs of residues to predict the native state conformation of a protein (Marks, et al. 2012; Sulkowska, et al. 2012; Sutto, et al. 2015), including repeat proteins (Espada, et al. 2015), as well as identify additional functionally relevant conformational states (Morcos, et al. 2013; Malinverni, et al. 2015; Sutto, et al. 2015) and multi-meric states (Schug, et al. 2009; Weigt, et al. 2009; Morcos, et al. 2011; dos Santos, et al. 2015; Malinverni, et al. 2015). Structural and coevolutionary information share complementary roles in the molecular simulations of proteins (See review: (Noel, et al. 2016)). The Potts model (Eq. 3) obtained from DCA has been related to the theory of evolutionary sequence selection (Morcos, et al. 2014) as well as mutational changes in protein stability (Lui and Tiana 2013; Morcos, et al. 2014; Contini and Tiana 2015). Additional work has applied DCA to protein folding to predict the effect of point mutations on the folding rate (Mallik, et al. 2016) as well as construct a statistical potential for native contacts in a structure-based model of a protein (Cheng, et al. 2016) to better capture the transition state ensemble.

DCA and inference methods have also been applied to study problems in systems biology, such as the identification of relevant protein-protein interactions in biological interaction networks (Procaccini, et al. 2011; Feinauer, et al. 2016). Recently, a number of studies have focused on inferring quantitative landscapes that capture the effects of mutations on biological phenotypes (Ferguson, et al. 2013; Cheng, et al. 2014; Figliuzzi, et al. 2016) by constructing models from sequence data (e.g., Eq. 3). Two separate studies, which focused on antibiotic drug resistance in *E. coli* (Figliuzzi, et al. 2016) and viral fitness of HIV-1 proteins (Ferguson, et al. 2013), respectively, inferred a Potts model (i.e., additive fitness landscape), *H*, and calculated mutational changes as Δ*H*. This approach is analogous to the approach adopted in this study. Likewise, a study examining the mutational effects on TCS phosphotransfer (Cheng, et al. 2014) constructed a mutational landscape from an information-based quantity. All of these approaches capture epistatic effects and rely on the accuracy of the inferred Potts model (Eq. 3) from sequence data.

### Mutational changes in coevolutionary landscape

For the concatenated sequences of HK (DHp) and RR (REC) (Fig. 1A), we infer a Potts model (Equation 3). We focus on a subset of parameters in our model consisting of the interprotein couplings, *J*_*ij*_, between positions in the DHp and REC domains (Fig. 1C) that are in close proximity in a representative structure of the TCS complex (Fig. 1B). All local fields terms, *h*_*i*_, are included to partially capture the fitness effects that give rise to the amino acid composition observed at each site. These considerations allow us to construct coevolutionary landscape for TCS, which is a negative, additive fitness landscape:

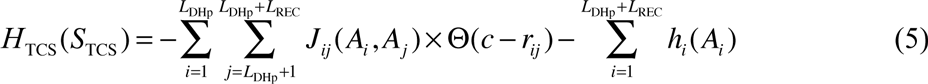

where *S*_TCS_ is the concatenated sequence of the DHp and REC domains, the double summation is taken over all interprotein statistical couplings between the DHp and REC domains, Θ is a Heaviside step function, *c* is the a cutoff distance of 16Å which was determined in a previous study (Morcos, et al. 2014), and *r*_*ij*_ is the minimum distance between residues *i* and *j* in the representative structure. Mutational changes in Eq. 5 are then computed as 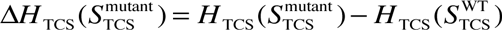. Hence, mutational changes in the signal transfer efficiency are approximated from mutational changes in the additive fitness.

An additional model is considered to analyze the epistatic effects of the 4-point mutations explored in experiment (Podgornaia and Laub 2015). While Eq. 5 naturally captures the epistatic effect of mutating a residue in the HK when the RR has also been mutated (or vice versa), it does not explicitly contain the statistical couplings that capture the epistatic effects of HK only mutations. Hence, the statistical couplings between the 4-mutational sites of the HK (Fig. 4C) are explicitly added to Eq. 5:

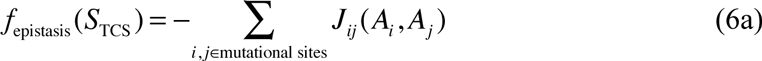

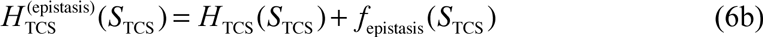

In Eq. 6a, the summation contains the 6 statistical couplings between the 4-mutational sites explored by Podgornaia and Laub, i.e., positions 14, 15, 18 and 19 in our MSA of TCS partners, which map to positions 254, 255, 258, and 259 of PhoQ.

### Zone Equilibration of Mutants (ZEMu) calculation

ZEMu consists of a multiscale minimization by dynamics, restricted to a flexibility zone of five residues about each substitution site (Dourado and Flores 2014), which is followed by a mutational change in stability using FoldX (Guerois, et al. 2002). ZEMu has been used to explain the mechanism of Parkinson’s disease associated mutations in Parkin (Caulfield, et al. 2014; Fiesel, et al. 2015). The minimization is done in MacroMoleculeBuilder (MMB), a general-purpose internal coordinate mechanics code also known for RNA folding (Flores and Altman 2010), homology modeling (Flores, et al. 2010), morphing (Tek, et al. 2016), and fitting to density maps (Flores 2014).

We use the Zone Equilibration of Mutants (ZEMu) (Dourado and Flores 2014) method to predict the mutational change in binding energy between PhoQ and PhoP. ZEMu first treats mutations as small perturbations on the structure by using molecular dynamics simulations (See Ref. (Dourado and Flores 2014) for full computational details) to equilibrate the local region around mutational sites. ZEMu can then estimate the binding affinity between the mutant-PhoQ/PhoP, 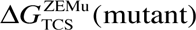, and the wild type-PhoQ/PhoP, 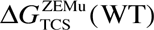, using the knowledge-based FoldX (Guerois, et al. 2002) potential. This allows for the calculation of the mutational change in binding affinity as: 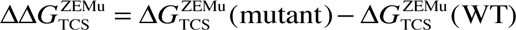.

ZEMu calculation was performed according to Ref: (Dourado and Flores 2014), with the following two differences. First, due to the large number of mutations we capped the computer time permitted to 3 core-hours per mutant, whereas in (Dourado and Flores 2014) the limit was 48 hours. This meant that of 122802 mutants attempted, 42923 completed within the time limit, whereas in (Dourado and Flores 2014), almost all mutants converged. The major reason for nonconvergence in the current work involved mutation to bulky or constrained residues. Steric clashes produced by such residues force the error-controlled integrator (Flores, et al. 2011) to take small time steps and hence use more computer time. Exemplifying this, the amino acids F, W and Y are the most common residues for non-converging mutations at positions 285 and 288 in PhoQ. The second difference was that we permitted flexibility in the neighborhood of all four possible mutation sites, even when not all of them were mutated, whereas in (Dourado and Flores 2014) only the mutated positions were treated as flexible. This allowed us to compare all of the mutational energies to a single wild type simulation, also performed with flexibility at all four sites.

Database of TCS partners, Potts model, and code for calculating Eq. 5 can be obtained from: http://utdallas.edu/~faruckm/PublicationDatasets.html

## Acknowledgments

We would like to thank Drs. Michele Di Pierro and Lena Simine for helpful comments.

## Author contributions

R.R.C., F.M., and J.N.O designed the research with the assistance of H.L., S.C.F., and O.N.; R.R.C and O.N. performed the research; R.R.C. and R.L.H. curated the protein databases that were used in our study; R.R.C., F.M., S.F. and O.N. wrote the paper.

